# Muscle-specific economy of force generation and efficiency of work production during human running

**DOI:** 10.1101/2021.02.18.431789

**Authors:** Sebastian Bohm, Falk Mersmann, Alessandro Santuz, Arno Schroll, Adamantios Arampatzis

## Abstract

Human running features a spring-like interaction of body and ground, enabled by elastic tendons that store mechanical energy and facilitate muscle operating conditions to minimize the metabolic cost. By experimentally assessing the operating conditions of two important muscles for running, the soleus and vastus lateralis, we investigated physiological mechanisms of muscle energy production and muscle force generation. Results showed that the soleus continuously shortened throughout the stance phase, operating as energy generator under conditions that were found to be optimal for work production: high force-length potential and enthalpy efficiency. The vastus lateralis promoted tendon energy storage and contracted nearly isometrically close to optimal length, resulting in a high force-length-velocity potential beneficial for economical force generation. The favorable operating conditions of both muscles were a result of an effective length and velocity-decoupling of fascicles and muscle-tendon unit mostly due to tendon compliance and, in the soleus, marginally by fascicle rotation.

## Introduction

During locomotion, muscles generate force and perform work in order to support and accelerate the body and the activation of the lower limb muscles accounts for most of the metabolic energy cost needed to walk or run [1–3]. Running is characterized by a spring-like interaction of the body with the ground, indicating temporally storage of kinetic and potential energy from the body in elastic elements, mainly tendons, as strain energy that can be recovered in the propulsive second half of the stance phase [3–5]. Storing mechanical energy in elastic tendons reduces the required energy production by the muscles through active shortening, which leads to lower metabolic energy cost [6–8] and a decrease in active muscle volume [4,9,10]. Thus, the consequence of this spring-like behavior is a reduction in the metabolic cost of running and an improvement in running economy.

At the muscle level, however, it has been shown that the triceps surae muscle group produces muscular work/energy during the stance phase of steady-state running [11]. The soleus is the largest muscle in this group [12] and does work by active shortening throughout the entire stance phase [13,14]. In the first part of the stance phase, the performed muscular work is stored in the Achilles tendon as elastic strain energy. During the later propulsion phase, the tendon strain energy recoil contributes to the muscular energy production, suggesting an energy amplification behavior [4] within the triceps surae muscle-tendon unit (MTU) during running. On the contrary, the vastus lateralis muscle (VL), as the main muscle of the quadriceps femoris muscle group [15], operates nearly isometrically despite a lengthening-shortening behavior of the VL MTU [16,17]. The almost isometric contraction suggests a negligible mechanical work production by the VL during running and a spring-like energy exchange between body and VL MTU, i.e. promoting energy conservation [3,4].

The triceps surae and the quadriceps muscle group are considered to be crucial for running performance [18,19]. The quadriceps femoris decelerates and supports the body early in stance while the triceps surae accounts for the propulsion later in the stance [18,20,21]. The soleus and VL, as the main muscles of both muscle groups, show marked differences in their morphological and architectural properties with shorter fascicles and higher pennation angles in the soleus [13,22] compared to VL [16,23]. Because of the long fascicles of the VL, a unit of force generated by this muscle is metabolically more expensive [10] compared to the soleus. Our previous findings [16] suggest that the VL operates at a high force-length-velocity potential (fraction of maximum force according to the force-length [24] and force-velocity [8] curves [13,16,25]) during running, which would indicate a fascicle contraction condition that could minimize the energetic cost of muscle force generation. The soleus muscle instead operates as an muscular energy generator through active shortening, which decreases the force-velocity potential [13,14] and may increase the energetic cost of muscle force generation, marking a trade-off between mechanical work production and metabolic expenses. When muscle fascicles shorten, the enthalpy efficiency [26] (or mechanical efficiency [27,28]) quantifies the fraction of ATP hydrolysis that is converted into mechanical work and depends on the shortening velocity, with a steep increase at low shortening velocities up to a maximum at around 20% of the maximum shortening velocity (V_max_) and a decrease thereafter [27–29]. Previous findings suggest that the soleus fascicles continuously shorten at a moderate velocity during the stance phase of running [13], covering a range that corresponds to a high efficiency. Therefore, the soleus muscle may operate at fascicle conditions that would be beneficial for economical work/energy production.

The muscle fascicle behavior is strongly influenced by the decoupling of the fascicles from the MTU excursions due to tendon elasticity and fascicle rotation [30–33]. The previously reported decoupling of the soleus muscle indicates that tendon elasticity and fascicle rotation affect the operating fascicle length and velocity during running [13,34], however their integration in the regulation of the efficiency-fascicle velocity dependency is unclear. Regarding the VL muscle, it was suggested that proximal muscles like the knee extensors feature shorter and less compliant tendons compared to the distal triceps surae muscles, thus limiting the decoupling between fascicles and MTU [35–37]. However, in our previous study, we found significantly smaller VL fascicle length changes compared to the VL MTU [16], indicating an important decoupling within the VL MTU due to tendon elasticity.

The purpose of this study was to assess the soleus and the VL fascicle behavior with regard to the operating force-length-velocity potential and enthalpy efficiency to investigate physiological mechanisms for muscle energy production and muscle force generation during running. We hypothesized that the soleus muscle as an energy generator operates at a high force-length potential and a high enthalpy efficiency, minimizing the metabolic cost of energy production. On the other hand, for the VL muscle that promotes energy conservation, we hypothesized a high force-length and a high force-velocity potential that would reduce the metabolic energy cost of muscle force generation. In order to investigate the regulation of the efficiency and force potentials, we further quantified the length and velocity decoupling of the fascicles from the MTU as well as the electromyographic (EMG) activation.

## Results

There were no significant differences in the anthropometric characteristics between groups (age p = 0.369, height p = 0.536, body mass p = 0.057). The experimentally assessed L_0_ of the soleus was on average 41.3 ± 5.2 mm and significantly shorter than L_0_ of the VL with 94.0 ± 11.6 mm (p < 0.001). The corresponding F_max_ of the soleus was 2887 ± 724 N, which was significantly lower compared to the 4990 ± 914 N of the VL (p < 0.001). Furthermore, the assessed V_max_ was 279 ± 35 mm/s for the soleus, significantly lower than the V_max_ of the VL with 1082 ± 133 mm/s (p < 0.001).

The stance and swing times during running were 304 ± 23 ms and 439 ± 26 ms for the soleus group and 290 ± 22 ms and 448 ± 30 ms for the VL group (p = 0.075, p = 0.369). The EMG comparison showed that the soleus was active throughout the entire stance phase of running while the VL was mainly active in the first part of the stance and with an earlier peak of activation (soleus 41 ± 5% of stance phase, VL 35 ± 4% of stance phase, p < 0.001, fig. 1). During the stance phase, the MTU of both muscles showed a lengthening-shortening behavior, but the VL MTU started to shorten earlier (soleus 59 ± 2% of stance phase, VL 50 ± 2% of stance phase, p < 0.001, fig. 1). The soleus and the VL fascicle length were clearly decoupled from the MTU length with smaller operating length ranges throughout the whole stance (fig. 1). The soleus fascicles operated at a length close to L_0_ at touchdown and then shortened continuously until the foot lift-off (0.994 to 0.752 L/L_0_, fig. 1). The operating length of the VL fascicles remained above L_0_ over the entire stance phase and was on average significantly longer compared to the soleus fascicles (soleus 0.899 ± 0.104 L/L_0_, VL 1.054 ± 0.082 L/L_0_, p < 0.001, fig. 1).

**Fig 1:**
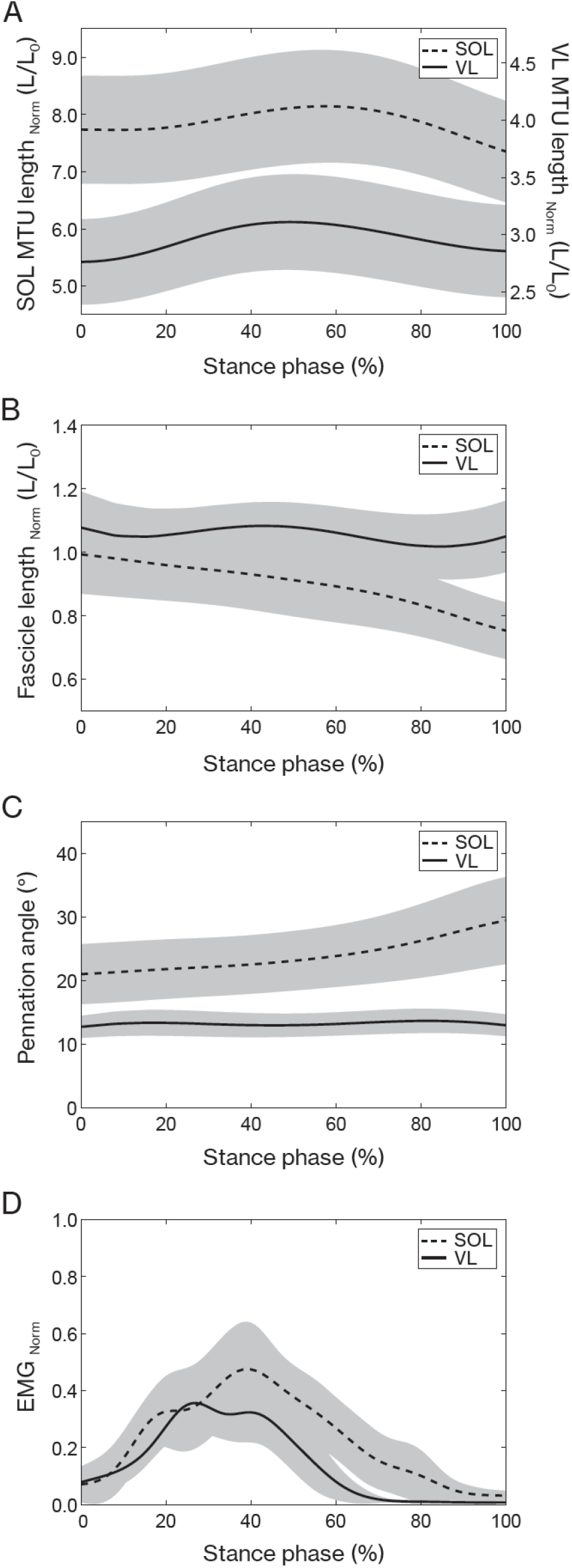
Soleus (SOL, n = 19) and vastus lateralis (VL, n = 14) muscle-tendon unit (MTU) length (A) and muscle fascicle length (normalized to optimal fascicle length L_0_, (B)), pennation angle (C) and electromyographic (EMG) activity (normalized to a maximum voluntary isometric contraction, (D)) during the stance phase of running (mean ± SD).

The stance phase-averaged force-length potential of both muscles was high and not significantly different (p = 0.689, fig. 2). The average pennation angle of the soleus was significantly greater than that of the VL (soleus 23.9 ± 5.1°, VL 13.3 ± 1.8°, p < 0.001) and increased continuously throughout stance, whereas it remained almost unchanged in the VL (fig. 1). The average operating velocity of the soleus fascicles was significantly higher compared to the VL (soleus 0.799 ± 0.260 L_0_ /s, VL 0.084 ± 0.258 L_0_ /s, p < 0.001), which showed an almost isometric contraction throughout stance. Consequently, the force-velocity potential (p < 0.001) and thus the overall force-length-velocity potential (p < 0.001) of the soleus was significantly lower compared to the VL during the stance phase (fig. 2).

**Fig 2:**
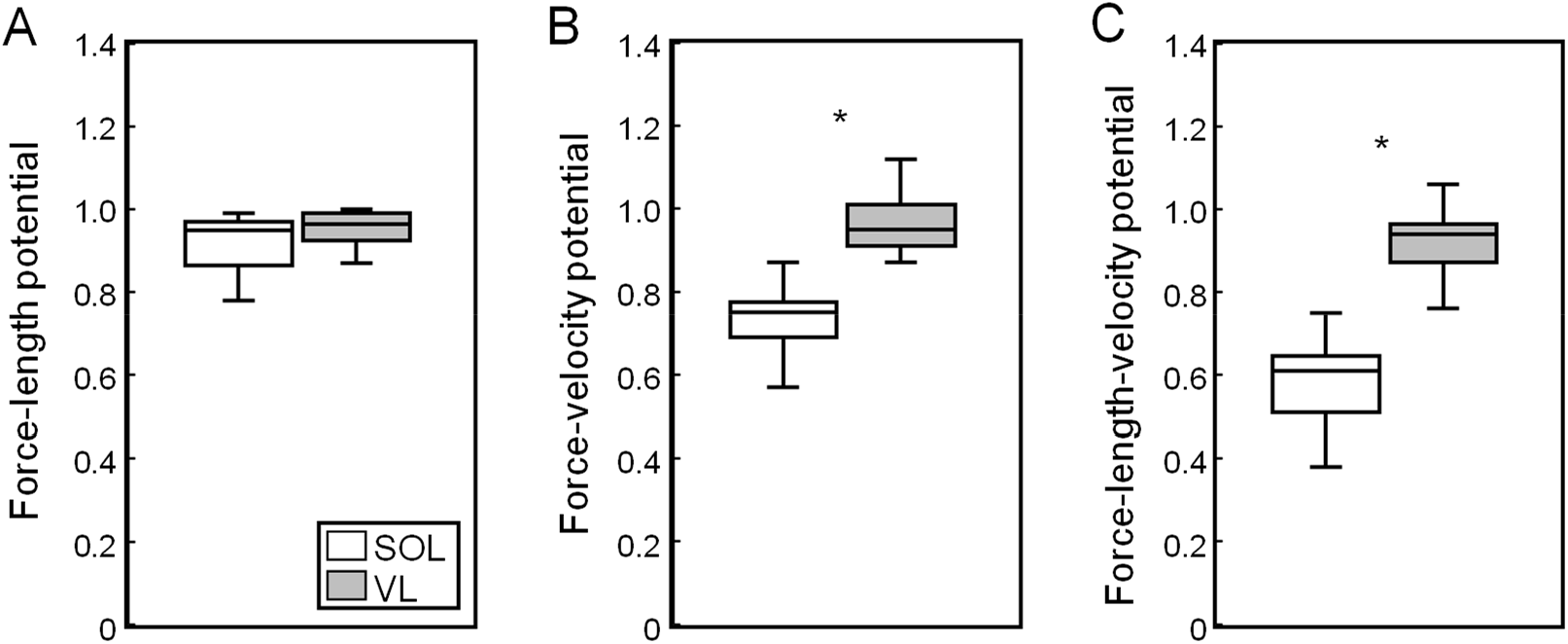
Soleus (SOL, n = 19) and vastus lateralis (VL, n = 14) force-length potential (A), force-velocity potential (B) and overall force-length-velocity potential (C) averaged over the stance phase of running. * significant difference between muscles (p < 0.05).

However, the higher shortening velocity of the soleus was close to the optimum one for maximum enthalpy efficiency, leading to a significantly higher enthalpy efficiency over the stance phase in comparison to the VL (p < 0.001, fig. 3).

**Fig 3:**
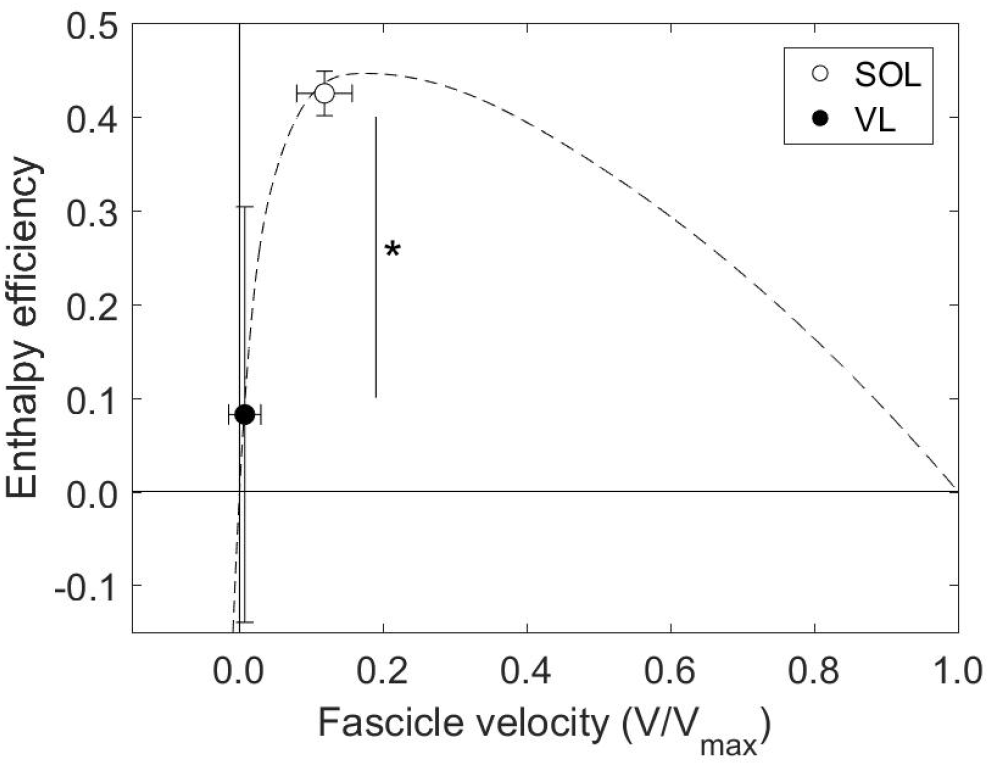
Soleus (SOL, n = 19) and vastus lateralis (VL, n = 14) enthalpy efficiency (mean ± SD) averaged over the stance phase of running onto the enthalpy efficiency-fascicle velocity relationship (dashed line). * significant difference between muscles (p < 0.05).

The fascicle, belly and MTU length changes throughout stance as well as the resulting velocity decoupling coefficients are illustrated in figure 4 for both muscles. There was a clear length- and velocity-decoupling of MTU and belly due to tendon compliance in both muscles (fig. 4). The SPM analysis revealed a significantly lower DC_Tendon_ of the soleus compared to the VL between 4 and 8% of stance phase (p = 0.032), since decoupling started later for the soleus. Between 20 and 57% of stance phase (p < 0.001) and between 65% of stance phase until lift-off, the soleus DC_Tendon_ was significantly higher than VL (p < 0.001, fig. 4). The DC_Tendon_ averaged over the stance phase of the soleus was also significantly greater (p < 0.001, tab. 1). Furthermore, the velocity-decoupling of belly and fascicles due to fascicle rotation progressively increased in the second part of the stance for the soleus but was negligible for the VL (fig. 4). The soleus DC_Belly_ was significantly higher from 33% of stance phase until lift-off compared to the VL as shown by the SPM analysis (p < 0.001, fig. 4) but also when averaged over the entire stance phase (p < 0.001, tab. 1). DC_Belly_ was markedly lower than DC_Tendon_, indicating that the tendon covered the majority of the overall decoupling in both muscles (fig. 4). Accordingly and similarly to DC_Tendon_, the SPM analysis for the overall decoupling of MTU and fascicles showed that DC_MTU_ of the soleus was significantly lower between 4 and 8% of stance phase (p = 0.032) and significantly higher from 20 to 57% of stance phase and from 65% of stance phase until lift-off compared to the VL (p < 0.001, fig. 4). The stance phase-averaged DC_MTU_ of the soleus was significantly greater compared to the VL as well (p <0.001, tab. 1).

**Fig 4:**
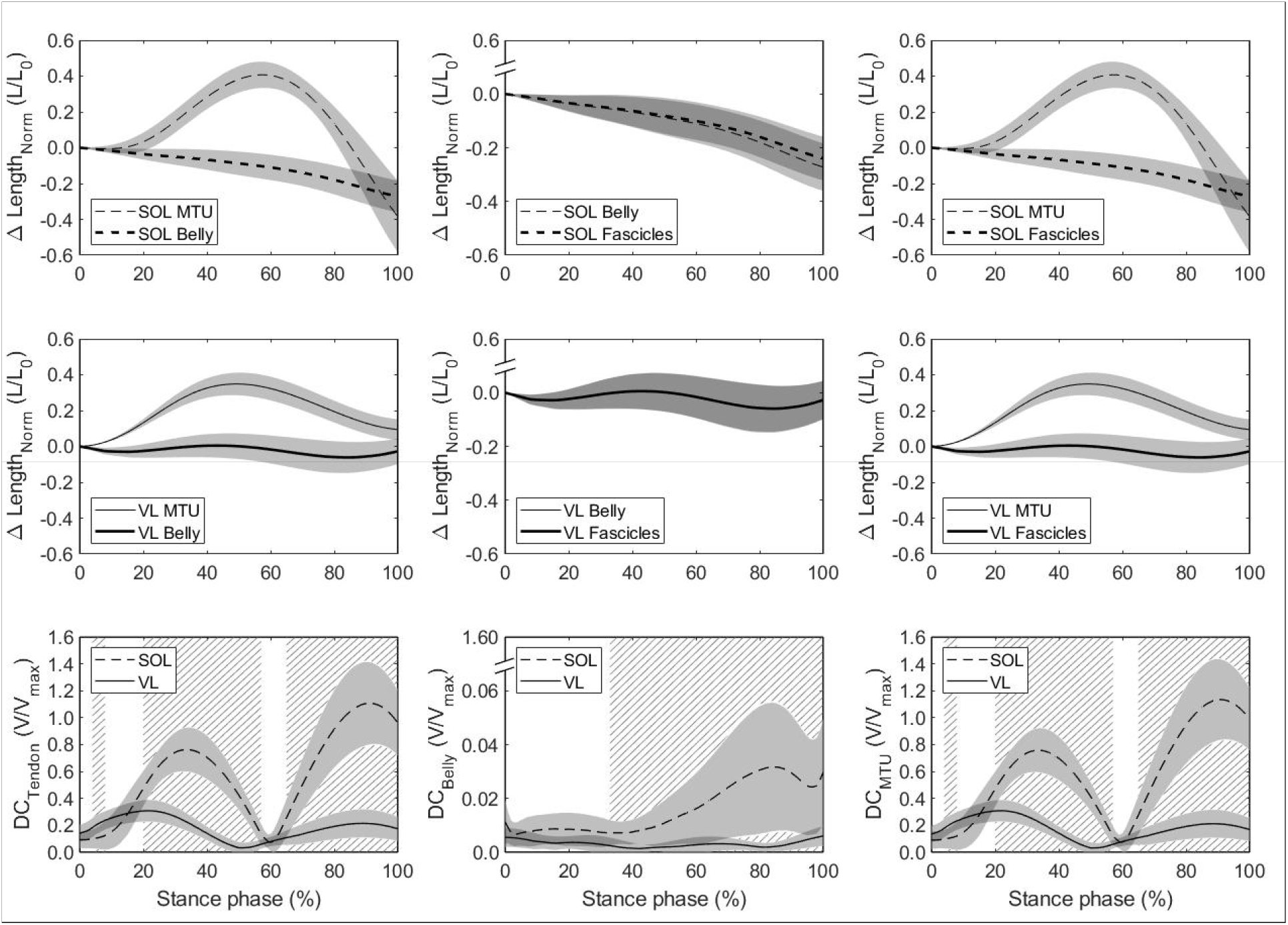
Soleus (SOL, n = 19, top row) and vastus lateralis (VL, n = 14, mid row) MTU vs. belly length changes (left), belly vs. fascicle length changes (mid) and MTU vs. fascicle length changes (right) over the stance phase of running with respect to the length at touchdown (0% stance phase). Differences between trajectories illustrate the length-decoupling due to tendon compliance, fascicle rotation and the overall decoupling, respectively. The bottom row shows the corresponding resulting velocity-decoupling coefficients (DC) as the absolute velocity differences between fascicles, belly and MTU normalized to the maximum shorting velocity (see methods). Intervals of stance with a significant difference between both muscles are illustrated as hatched areas (p < 0.05).

## Discussion

We mapped the operating length and velocity of the soleus and the VL fascicles during running onto the individual force-length, force-velocity and enthalpy efficiency-velocity curves in order to investigate physiological mechanisms for muscle force generation and muscle energy production in the two muscles. The soleus continuously shortened throughout the stance phase and produced muscular work at a shortening velocity close to the enthalpy efficiency optimum. VL operated with smaller length changes, almost isometrically, resulting in a high force-velocity potential beneficial for economic force generation. Both muscles operated close to L_0_, i.e. at a high force-length potential. Tendon compliance covered the majority of the overall decoupling of MTU and fascicles in both muscles, enabling favorable conditions for muscle force or muscle work production. Only in the soleus muscle, fascicle rotation contributed to the overall decoupling, indicating an additional, yet comparatively minor, effect on the fascicle dynamics during locomotion.

The triceps surae and quadriceps muscle groups are the main actuators for locomotion and thus responsible for a great portion of the metabolic energy cost of running [18,36,38,39]. While the quadriceps mainly decelerates and supports body mass in the early stance phase, the triceps surae contributes to the acceleration of the center of mass during the second part of the stance phase [18,20]. The soleus is the largest muscle of the triceps surae [12] and the VL of the quadriceps [15] and thus both muscles are important contributors to the running movement. We found that the soleus actively shortened throughout the entire stance phase, indicating continuous work/energy production. The average velocity at which the soleus shortened was very close to the optimal velocity for maximal enthalpy efficiency. Enthalpy efficiency quantifies the fraction of chemical energy from ATP hydrolysis that is converted into mechanical muscular work [26,27] with a peak at around 20% of V_max_ [27,29]. Consequently, the mechanical work performed by the soleus muscle, being essential during running [18,40–42] and high enough in magnitude to significantly influence the overall metabolic energy cost of locomotion [13,43,44], was generated at a high enthalpy efficiency (94% of maximum efficiency). Considering that also the soleus force-length potential was close to the maximum (0.92) and that a high potential may decrease the active muscle volume for a given muscle force [9,10,43], our results provide evidence of a cumulative contribution of two different mechanisms (high force-length potential and high enthalpy efficiency) to an advantageous muscular energy production of the soleus during running. The VL was mainly active in the first part of the stance phase and its fascicles operated with very small length changes, i.e. almost isometrically, confirming earlier reports [16,17]. This indicates that the VL dissipates and/or produces negligible amounts of mechanical energy during running, yet generating force for the deceleration and support of the body mass. The found decoupling of the VL MTU and fascicles showed that the deceleration of the body mass in the early stance phase was not a result of an energy dissipation by the contractile element (active stretch) but rather an energy absorption by the tendinous tissue. Tendons feature small damping characteristics resulting in a hysteresis of only 10% [45,46] and, therefore, the main part of the absorbed energy of the body’s deceleration is stored as elastic tendon strain energy, which is then returned later in the second part of the stance phase. The high force-length (0.93) and force-velocity (0.90) potential of the VL muscle throughout stance indicates an energy exchange within the VL MTU under almost optimal conditions for muscle force generation during running. Operating at high potentials reduces the active muscle volume for a given force [9,10] and thus the metabolic energy cost of muscle force generation.

By actively shortening, the soleus delivered energy during the entire stance phase to the skeleton, providing the main muscular work required for running. On the other side, the contractile elements of the VL muscle did not contribute to the required muscular work and operated in concert with the elastic tendon in favor of energy storage [4]. Our findings showed that, although the human body interacts with the ground in a spring-like manner during steady-state running to conserve mechanical energy [3,4], there are indeed muscles that operate as energy generators, like the soleus, and others that promote energy conservations, like the VL. Further, our results indicate that the fascicle operating length and velocity of the soleus muscle, the main energy generator, is optimized for high enthalpy efficiency, while those of the VL muscle, that promote energy conservation, for a high potential of force generation. The consequence of the active shortening of the soleus muscle for work production is a decrease of the force-velocity potential during the stance phase, which may increase the active muscle volume and shortening-related cost [6–8]. However, the soleus muscle features shorter fascicles (L_0_ = 41 mm) compared to the VL muscle (L_0_ = 94 mm) and, for this reason, a given force generated by the soleus is energetically less expensive [10]. The specific morphology of the soleus muscle certainly compensates for the reductions of the force-velocity potential and provides advantages for its function as energy generator during submaximal steady-state running. Furthermore, operating around the “sweet spot” of the shortening velocity for high enthalpy efficiency facilitates the economical muscular work production, while either a too high or a too low shortening velocity would be disadvantageous.

The almost optimal conditions for muscular work production and muscle force generation of the soleus and VL were a result of an effective decoupling between MTU and fascicle length that was regulated by an appropriate muscle activation. For the soleus, the activation level increased in the first part of stance phase, contracting the muscle while the MTU increased in length. This activation pattern not only prevented the muscle to be stretched but also induced continuous shortening around the plateau of the force-length curve at a high enthalpy efficiency. The respective high DC_Tendon_ further indicates that a part of the body’s mechanical energy was stored as strain energy in the Achilles tendon in addition to the generated work by fascicle shortening. During MTU shortening (propulsion phase), the soleus EMG activation decreased and the tendon recoiled, enabling the high shortening velocities of the MTU while maintaining the fascicle operating conditions close to the efficiency optimum. The simultaneous release of the stored strain energy from the tendon further added to the ongoing muscle work production, i.e. energy amplification. The VL muscle showed higher levels of activation during the initial part of the stance phase and earlier deactivation than soleus. The timing and level of activation regulated the decoupling within the VL MTU during the body mass deceleration in a magnitude that the lengthening and shorting of the MTU was fully accomplished by the tendinous tissue. Consequently, the VL fascicles operated at a high force-length-velocity potential and the body’s energy was conserved within the MTU. Although being substantial for soleus and VL, the SPM analysis revealed higher values of DC_Tendon_ for soleus during the major part of the stance phase (average value for soleus 0.57 V/V_max_ and VL 0.18 V/V_max_), indicating a greater decoupling within the soleus MTU compared to the VL MTU. In the soleus muscle, fascicle rotation (changes in pennation angle) had an additional effect on the overall decoupling between MTU and fascicles. The results showed an increase in DC_Belly_ in the second part of the stance phase where the soleus belly velocity was high during the MTU shortening. However, the decoupling by the fascicle rotation was considerable smaller compared to the tendon decoupling. Over the stance phase, belly and tendon decoupling were 1.6 %V_max_ and 57 %V_max_ and during the MTU shortening phase 2.6 %V_max_ and 72 %V_max_ respectively, suggesting a rather minor functional role of fascicle rotation during submaximal running. In the VL, fascicle rotation was virtually absent and consequently DC_Belly_ values showed no relevant decoupling effect at all.

In conclusion, our results showed that during the stance phase of steady-state running, when the human body interacts with the environment in a spring-like manner, the soleus muscle acts as energy generator and the VL muscle as energy conservator. Furthermore, our findings provide evidence that the soleus operates under conditions optimal for muscular energy production (i.e. high force-length potential and high enthalpy efficiency) and the VL under conditions optimal for muscle force generation (i.e. high force-length and high force-velocity potential).

## Materials and methods

### Participants and experimental design

Thirty-three physically active adults were included in the present investigation. None of the participants reported any history of neuromuscular or skeletal impairments in the six months prior to the recordings. The ethics committee of the university approved the study (EA2/076/15) and the participants gave written informed consent in accordance with the Declaration of Helsinki. From the right leg, either the soleus (n = 19, 29 ± 6 yrs., 177 ± 9 cm, 69 ± 9 kg, 7 females) or vastus lateralis (n = 14, age 28 ± 4 yrs., height 179 ± 7 cm, body mass 75 ± 8 kg, 3 females) muscle fascicle length, fascicle pennation angle and EMG activity were recorded during running on a treadmill at 2.5 m/s. Corresponding MTU lengths were calculated from the kinematic data and individually measured tendon lever arms. We further assessed the soleus and VL force-fascicle length and force-fascicle velocity relationship to calculate the force-length and force-velocity potential of the soleus and the VL muscle fascicles during running. The operating fascicle velocity was additionally mapped on the enthalpy efficiency-velocity relationship to assess the enthalpy efficiency of both muscles. The contribution of the decoupling of the fascicle length and velocity from the MTU to the operating force potential and enthalpy efficiency at the level of tendon and muscle belly during running was examined for both muscles as well.

### Joint kinematics, fascicle behavior and electromyographic activity during running

After a familiarization phase, a four-minute running trial on a treadmill (soleus: h/p cosmos mercury, Isny, Germany; VL: Daum electronic, ergo_run premium8, Fürth, Germany) was performed and kinematics of the right leg were captured by a Vicon motion capture system (version 1.8, Vicon Motion Systems, Oxford, UK, 250 Hz) using an anatomically-referenced reflective marker setup (greater trochanter, lateral femoral epicondyle and malleolus, fifth metatarsal and tuber calcanei). The kinematic data were used to determine the touchdown of the foot and the toe-off as consecutive minima in knee joint angle over time [47]. Furthermore, the kinematics of the ankle and knee joint served to calculate the MTU length change of the soleus and VL during running, as the product of ankle joint angle changes and Achilles tendon lever arm as well as knee joint angle changes and patellar tendon lever arm [48], respectively. We used the ultrasound-based tendon-excursion method for the Achilles tendon lever arm determination [49]. The patellar tendon lever arm was measured using magnetic resonance imaging in fully extended knee joint position and calculated as a function of the knee joint angle change using the data by Herzog & Read [50] (for a detailed description of both tendon lever arm measurements see [13,14,16]). The initial soleus and VL MTU length was calculated based on the regression equation provided by Hawkins & Hull [51] at neutral ankle joint angle for the soleus MTU and at touchdown for the VL MTU. During the running trial, ultrasound images of either the soleus or VL muscle fascicles were recorded synchronously to the kinematic data (soleus: Aloka Prosound Alpha 7, Hitachi, Tokyo, Japan, 6 cm linear array probe, UST-5713T, 13.3 MHz, 146 Hz; VL: My Lab60, Esaote, Genova, Italy, 10 cm linear array probe LA923, 10 MHz, 43 Hz). The ultrasound probe was mounted over the medial aspect of the soleus muscle belly or on the VL muscle belly (≈50% of femur length) using a custom anti-skid neoprene-plastic cast. The fascicle length was post-processed from the ultrasound images using a self-developed semi-automatic tracking algorithm [23] that calculated a representative reference fascicle on the basis of multiple muscle fascicle portions identified from the entire displayed muscle (for details see [16,23], fig. 5). Visual inspection of each image was conducted and corrections were made if necessary. At least nine steps were analyzed for each participant and then averaged [16,52]. The pennation angle was calculated as the angle between the deeper aponeurosis and the reference fascicle (fig. 5). The length changes of the muscle belly of soleus and VL were calculated as the differences of consecutive products of fascicle length and the respective cosine of the pennation angle [53]. Note that this does not give the length of the entire soleus or VL muscle belly but rather the projection of the instant fascicle length onto the plane of the MTU, which can be used to calculate the changes of the belly length [13]. The velocities of fascicles, belly and MTU were calculated as the first derivative of the lengths over time. Surface EMG of the VL and the soleus was measured by means of a wireless EMG system (Myon m320RX, Myon AG, Baar, Switzerland, 1000 Hz). A fourth-order high-pass Butterworth filter with 50 Hz cut-off frequency, a full-wave rectification and then a low-pass filter with 20 Hz cut-off frequency were applied to the raw EMG data. The EMG activity was averaged over the same steps that were analyzed for the soleus parameters and for the VL over 10 running steps. EMG values were then normalized for each participant to the maximum obtained during the individual MVCs.

**Fig 5.**
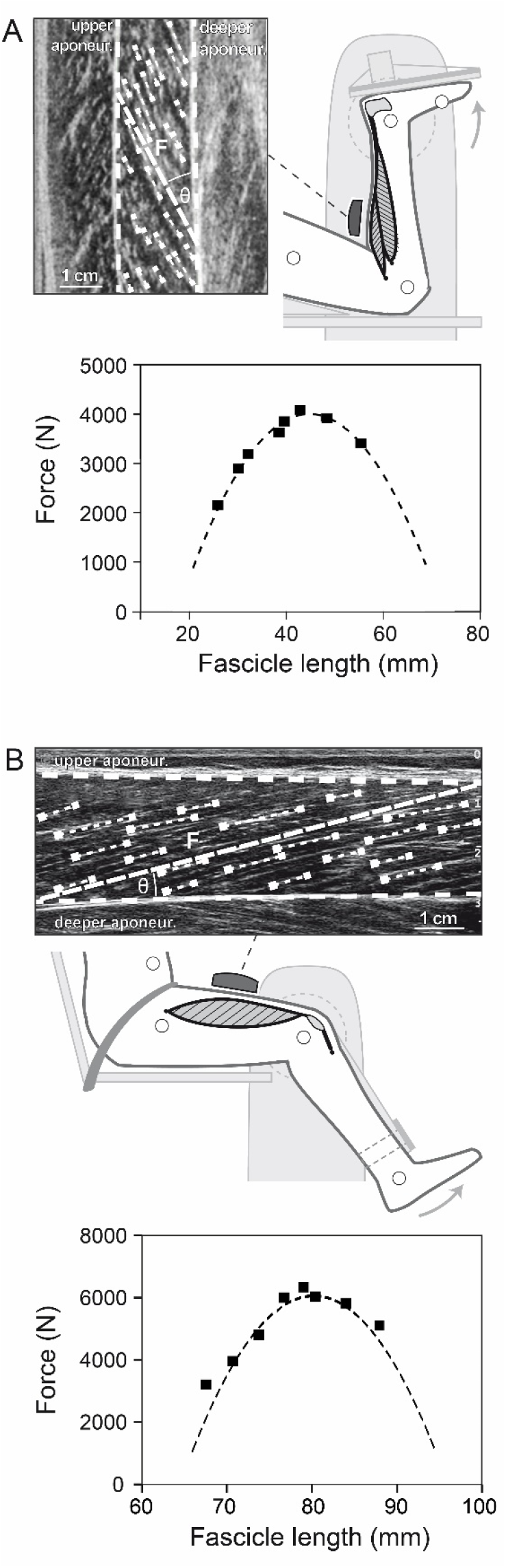
Experimental setup for the determination of the soleus (A) and vastus lateralis (VL, B) force-fascicle length relationship. Maximum isometric plantar flexions (MVC) at eight different joint angles was performed on a dynamometer. During the MVCs, ultrasound images of the soleus and VL were recorded and a representative muscle fascicle length (F) was calculated based on multiple fascicle portions (short dashed lines). Accordingly, an individual force-fascicle length relationship for the soleus and VL muscle was derived from the MVCs (squares) by means of a second-order polynomial fit (dashed line, bottom graphs).

### Assessment of the force-length, force-velocity and enthalpy efficiency-velocity relationship

To determine the soleus and the VL force-length relationship, eight maximum voluntary plantar flexion or knee extension contractions (MVCs) in different joint angles were performed with the right leg on an isokinetic dynamometer (Biodex Medical, Syst. 3, Inc., Shirley, NY), following a standardized warm-up [13,16,25] (fig. 5). For the plantar flexion MVCs, the participants were placed in prone position with the knee in fixed flexed position (∼120°) to restrict the contribution of the bi-articular m. gastrocnemius to the plantar flexion moment [54] and the joint angles were set in a randomized equally-distributed order ranging from 10° plantar flexion to the individual maximum dorsiflexion angle. Regarding the knee extensions, participants were seated with a hip joint angle of 85° to reduce the contribution of the bi-articular m. rectus femoris [55], while the knee joint angle ranged between 20° to 90° knee joint angle (0° = knee extended) in randomly ordered 10° intervals. The resultant moments at the ankle and knee joint were calculated under consideration of the effects of gravitational and passive moments and any misalignment between joint axis and dynamometer axis using an established inverse dynamics approach [56,57]. The required kinematic data were recorded during the MVCs based on anatomically referenced reflective markers (medial and lateral malleoli and epicondyle, calcaneal tuberosity, second metatarsal and greater trochanter) by a Vicon motion capture system (250 Hz). Furthermore, the contribution of the antagonistic moment produced by tibialis anterior during the plantar flexion MVCs or by the hamstring muscles during the knee extension MVCs was taken into account by means of an EMG-based method according to Mademli et al. [58]. The force applied to the Achilles tendon or patellar tendon during the plantar flexion or knee extension MVCs was calculated as quotient of the joint moment and individual tendon lever arm, respectively. The soleus or the VL fascicle behavior during the MVCs was synchronously captured by ultrasonography and fascicle length was determined using the same methodology described above (fig. 5). Accordingly, an individual force-fascicle length relationship was calculated for soleus or VL by means of a second-order polynomial fit and the maximum muscle force applied to the tendon (F_max_) and optimal fascicle length for force generation (L_0_) was derived, respectively (fig. 5).

The force-velocity relationship of the soleus and the VL muscle was further assessed using the classical Hill equation [8] and the muscle-specific V_max_ and constants of a_rel_ and b_rel_. For V_max_ we took values of human soleus and VL type 1 and 2 fibers measured in vitro at 15°C reported by Luden et al. [59]. The values were then adjusted [60] for physiological temperature conditions (37 °C) and an average fiber type distribution of the human soleus (type 1 fibers: 81%, type 2: 19%) and VL muscle (type 1 fibers: 37%, type 2: 63%) reported in literature [59,61–63] was the basis to derive a representative value of V_max_. For the soleus muscle under the in vivo condition, V_max_ was calculated as 6.77 L_0_ /s and for the VL as 11.51 L_0_ /s. For L_0_ we then referred to the individually measured optimal fascicle length (described above, fig. 5). The constant a_rel_ was calculated as 0.1+0.4FT, where FT is the fast twitch fiber type percentage, which then equals to 0.175 for the soleus and 0.351 for the VL [64,65]. The product of a_rel_ and V_max_ gives the constant b_rel_ as 1.182 for the soleus and 4.042 for the VL [66]. Based on the assessed force-length and force-velocity relationships, we calculated the individual force-length and force-velocity potential of both muscles as a function of the fascicle operating length and velocity during the stance phase of running. The product of both potentials then gives the overall force-length-velocity potential.

Furthermore, we determined the enthalpy efficiency-velocity relationship for the soleus and the VL muscle fascicles in order to calculate the enthalpy efficiency of both muscles as a function of the fascicle operating velocity during running. For this purpose, we used the experimental efficiency values provided by the paper of Hill 1964 in table 1 for a/P_0_ = 0.25 [27]. Because the effect of differences in a/P_0_ on the shape of the curve is negligible [27], we used the same values for both muscles. By means of the classical Hill equation [8], we then transposed the original efficiency values that were presented as a function of relative load (relative to maximum tension) to shortening velocity (normalized to V_max_). The values of enthalpy efficiency and shortening velocity were then fitted using a cubic spline, giving the right-skewed parabolic-shaped curve with a peak efficiency of 0.45 at a velocity of 0.18 V/V_max_. The resulting function was then used to calculate the enthalpy efficiency of the soleus and the VL during running based on the average value of the fascicle velocity over stance, accordingly.

**Table 1:** Average tendon (DC_Tendon_), belly (DC_Belly_) and muscle-tendon unit (DC_MTU_) decoupling coefficients for the soleus and vastus lateralis (VL) muscles during the stance phase of running (mean ± SD).

|  | Soleus (n=19) | VL (n=14) |
| --- | --- | --- |
| $DC_{Tendon} (V/V_{max})$ | $0.567 \pm 0.128$ | $0.180 \pm 0.053^*$ |
| $DC_{Belly} (V/V_{max})$ | $0.016 \pm 0.008$ | $0.003 \pm 0.002^*$ |
| $DC_{MTU} (V/V_{max})$ | $0.574 \pm 0.127$ | $0.179 \pm 0.014^*$ |

\* Statistically significant difference between the two muscles ( $p < 0.05$ )

### Assessment of decoupling within the muscle-tendon unit

To quantify the decoupling of fascicle, belly and MTU velocities over the time course of stance we calculated a decoupling coefficient to account for the tendon compliance (DC_Tendon_, equation 1), fascicle rotation (DC_Belly_, equation 2) as well as for the overall decoupling of MTU and fascicle velocities that includes both components (DC_MTU_, equation 3).

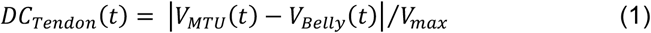

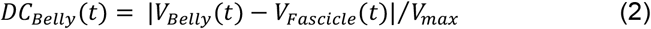

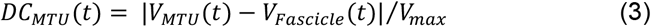

*V(t)* is the velocity at each percentage of the stance phase (i.e. t = 0, 1, …, 100 %stance). We introduced these new decoupling coefficients because previously suggested decoupling ratios (i.e. tendon gearing = V_MTU_ /V_Belly_, belly gearing (or architectural gear ratio) = V_Belly_ /V_Fascicle_, MTU gearing = V_MTU_ /V_Fascicle_ [30,31]) may feature limitations for the application under in vivo conditions, i.e. considering that muscle belly and fascicle velocities may be very close to or even zero during functional tasks as walking and running [13,16], which results in non-physiological gear ratios.

### Statistics

A t-test for independent samples was used to test for group differences in anthropometric characteristics, temporal gait parameters and differences between the soleus and the VL fascicle belly, MTU and EMG parameters. The Mann-Whitney U test was applied in case the assumption of normal distribution, tested by the Kolmogorov-Smirnov test with Lilliefors correction, was not satisfied. The level of significance was set to α = 0.05 and the statistical analyses were performed using SPSS (IBM Corp., version 22, NY, US). Furthermore, statistical parametric mapping (SPM, independent sample t-test, α = 0.05) was used to test for differences between the DC_Tendon_, DC_Belly_ and DC_MTU_ of the soleus and the VL throughout the stance phase of running. SPM was conducted using the software package spm1D (version 0.4, www.spm1d.org) [67].

## Authors’ contributions

S.B., F.M., A.S., A.S. and A.A. designed research. S.B., F.M. and A.S. performed research. S.B. analysed data. S.B. and A.A. drafted the manuscript. F.M., A.S. and A.S. made important intellectual contributions during revision.

